# SARS-CoV-2 neutralizing antibody specificities differ dramatically between recently infected infants and immune-imprinted individuals

**DOI:** 10.1101/2025.01.17.633612

**Authors:** Bernadeta Dadonaite, Allison R. Burrell, Jenni Logue, Helen Y. Chu, Daniel C. Payne, David B. Haslam, Mary A. Staat, Jesse D. Bloom

## Abstract

The immune response to viral infection is shaped by past exposures to related virus strains, a phenomenon known as imprinting. For SARS-CoV-2, much of the population has been imprinted by a viral spike from an early strain, either through vaccination or infection during the early stages of the COVID-19 pandemic. As a consequence of this imprinting, infection with more recent SARS-CoV-2 strains primarily boosts cross-reactive antibodies elicited by the imprinting strain. Here we compare the neutralizing antibody specificities of imprinted individuals versus infants infected with a recent strain. Specifically, we use pseudovirus-based deep mutational scanning to measure how spike mutations affect neutralization by the serum antibodies of adults and children imprinted by the original vaccine versus infants with a primary infection by a XBB* variant. While the serum neutralizing activity of the imprinted individuals primarily targets the spike receptor-binding domain (RBD), serum neutralizing activity of infants only infected with XBB* mostly targets the spike N-terminal domain (NTD). In these infants, secondary exposure to the XBB* spike via vaccination shifts more of the neutralizing activity towards the RBD, although the specific RBD sites targeted are different than for imprinted adults. The dramatic differences in neutralization specificities among individuals with different exposure histories likely impact SARS-CoV-2 evolution.

## Introduction

Human antibody responses to viral infections are influenced by infection history (1). For SARS-CoV-2, much of the global population first encountered the virus through infection or vaccination by strains with spikes very similar to the early Wuhan-Hu-1 strain (2,3). This initial exposure has shaped immune responses to subsequent infections and vaccinations, a phenomenon known as imprinting (4,5). While imprinting primes the immune system to respond effectively to closely related strains, it can hinder the generation of new B-cells specific to more diverged SARS-CoV-2 strains. For example, individuals imprinted with the early Wuhan-Hu-1 strain primarily activate cross-reactive B-cells upon infection with newer variants (3,6,7). These B-cells can produce antibodies that neutralize both the original imprinting strain and the newer variant, but the overall magnitude of the neutralizing response to the new variant is often lower than the original response to the imprinting strain (2,8–10).

Most previous studies of the specificities of SARS-CoV-2 neutralizing antibodies have focused on individuals imprinted with early pre-Omicron variants (2,10,11) but individuals with their first exposure to post-Omicron strains have not been as widely studied. In addition, because mutations in the spike’s receptor binding domain (RBD) drive much of the antigenic evolution in SARS-CoV-2 (12–14), many studies have focused primarily on analyzing antibodies that bind to the RBD (6,15–17). However, during SARS-CoV-2 evolution in humans, the N-terminal domain (NTD) has also undergone extensive antigenic changes (18–20).

Here we compare the neutralizing antibody responses between humans imprinted by early SARS-CoV-2 vaccination versus infection with a more recent variant. To do this, we use pseudovirus deep mutational scanning (21,22) to measure the effects of mutations to the XBB.1.5 spike on neutralization by sera from adults and children first vaccinated with the Wuhan-Hu-1 strain versus infants whose initial exposure was infection with the XBB* variants that were dominant in 2023. We find that sera from primary XBB* infections exhibit markedly different neutralizing specificities compared to Wuhan-Hu-1-imprinted sera. Specifically, neutralizing activity in primary XBB* infection sera predominantly targets the NTD, while neutralization by sera from imprinted individuals focuses on the RBD. However, additional exposures to the XBB* spike broaden the neutralizing response to strongly target both the NTD and RBD, correlating with increased sera potency. Additionally, we identify mutations with contrasting effects depending on imprinting history—some mutations that reduce neutralization by Wuhan-Hu-1-imprinted sera actually enhance neutralization by XBB*-imprinted sera and vice-versa. These findings underscore how diverse infection histories shape neutralizing antibody responses to spike, and in turn drive the evolution of new SARS-CoV-2 variants.

## Results

### Sera from humans imprinted with the early Wuhan-Hu-1 strain or a later XBB* strain

To examine the effects of imprinting on the specificity of polyclonal serum neutralizing antibody activity targeting the SARS-CoV-2 spike, we assembled several sets of human sera representing different exposure histories (**Fig. 1A**, **Table S1**). Six sera were collected from infants after their first infection with a XBB* variant in 2023; we refer to these as *primary XBB* infection infant sera*. These infants were all at least six months old at the time of their first infection which is old enough that maternal antibodies to SARS-CoV-2 should have largely waned, which was validated by ELISAs on pre-infection serum samples (**Fig. S1**). Two of these infants later received the XBB.1.5 variant vaccine, recommended in September 2023 by the US Advisory Committee on Immunization Practices for all persons ≥6 months (23); we refer to sera collected after these vaccinations as *XBB* infected and vaccinated infant sera*. We previously published antigenic maps of sera from adults who were vaccinated multiple times with original Wuhan-Hu-1 spike-containing vaccines starting in late 2020 and subsequently exposed to different SARS-CoV-2 variants (21). Based on the date of their last infection in 2023 or sequencing of the infecting strain, these adults were either confirmed or suspected to have last been exposed to the XBB* variant and are therefore referred to here as *Wuhan-Hu-1-imprinted and XBB* infected adult sera*. Similarly, *Wuhan-Hu-1 imprinted and XBB* infected children sera* were collected from children who had received at least two doses of the original Wuhan-Hu-1 spike-containing vaccine and were subsequently infected with the XBB* variant. The infant and children sera were from the IMPRINT cohort at Cincinnati Children’s Hospital Medical Center, which collects weekly nasal swabs from participants starting in the second week of life, allowing for detailed infection history tracking and sequence validation for variant infections; the adult sera were from the HAARVI cohort in Seattle (**Table S1**).

**Figure 1.**
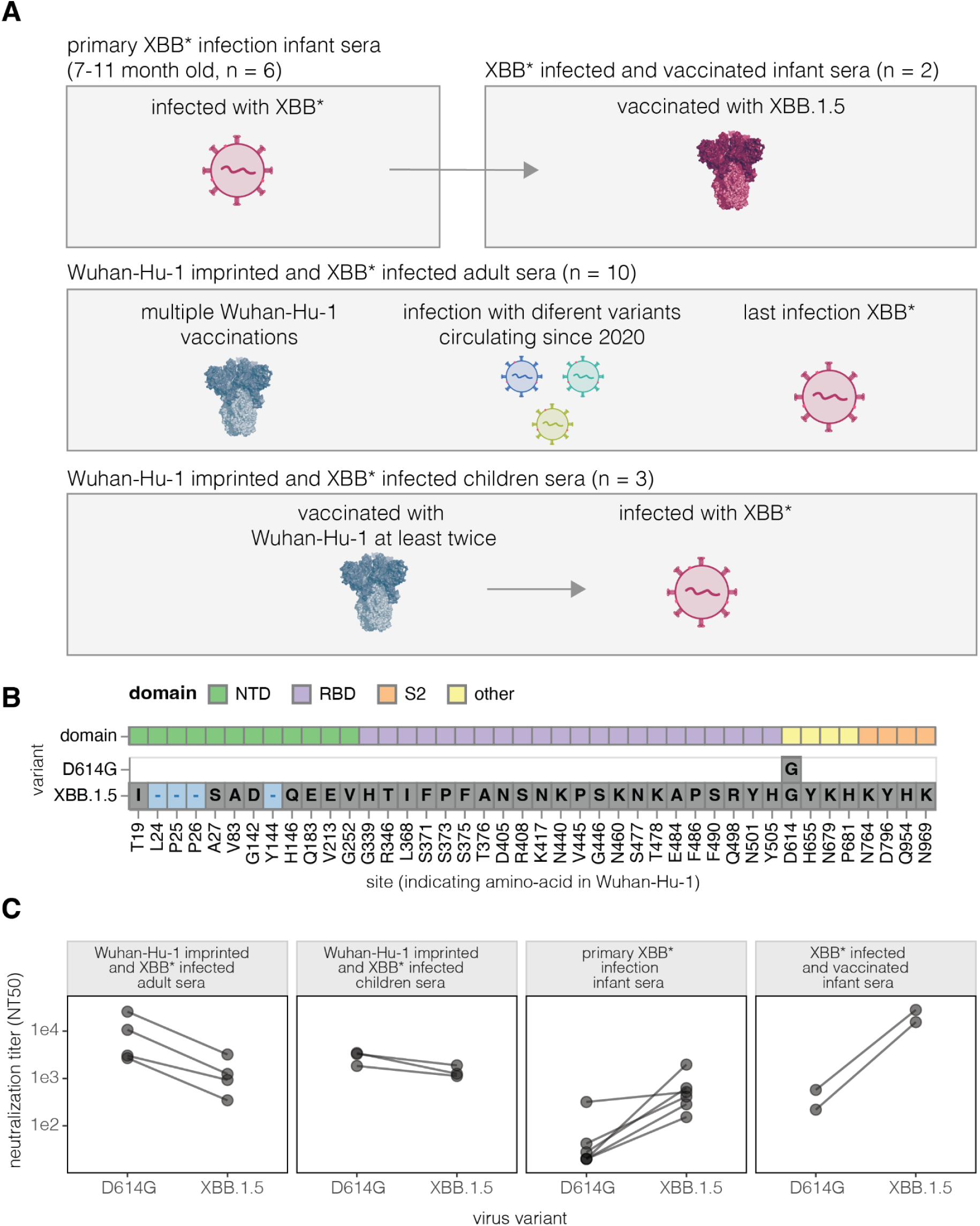
Sera from humans imprinted with Wuhan-Hu-1 or XBB* variants. **A.** Sets of sera used in this study. For detailed description of individual sera see **Table S1**. **B.** Amino-acid mutations in the D614G and XBB.1.5 spike relative to the original Wuhan-Hu-1 spike. **C** Neutralization of pseudoviruses expressing the spike from D614G or XBB.1.5 by each set of sera. For the *Wuhan-Hu-1 imprinted and XBB* infected adult* set, neutralization assays were performed on four representative sera.

The Wuhan-Hu-1 and XBB.1.5 spikes differ by 42 amino-acid mutations, with the majority of these mutations in the RBD and the next largest number of mutations in the NTD (**Fig. 1B**). The Wuhan-Hu-1 spike matches that of the earliest known SARS-CoV-2 variant from 2019, whereas the XBB.1.5 variant was dominant in mid-2023. Note that we use “XBB*” to refer to the larger group of XBB-related variants that circulated in 2023. For the experiments in this paper, we perform assays on the D614G rather than Wuhan-Hu-1 spike; the D614G variant became dominant in the first half of 2020 and contains only the D614G substitution relative to Wuhan-Hu-1 (24). We use the D614G spike rather than the Wuhan-Hu-1 spike as it is well known to yield better pseudovirus titers without appreciably affecting the binding of neutralizing antibodies (24,25).

The sera from both adults and children imprinted with Wuhan-Hu-1 and subsequently infected by XBB* variants had high neutralizing titers against pseudoviruses with either the early D614G spike or the XBB.1.5 spike (**Fig. 1C**). For these sera, the titers were higher against the D614G than XBB.1.5 spike, in agreement with numerous prior studies showing that neutralizing titers are typically highest against the imprinting strain (2,8,9,26). By contrast, most sera from infants with primary XBB* infection showed little neutralization of the D614G spike but had high titers to the XBB.1.5 variant (**Fig. 1C**), consistent with the fact these sera were elicited by XBB* strains. Two of the infants infected with a XBB* variant were subsequently vaccinated with the XBB.1.5 spike, after which their titers against XBB.1.5 increased to levels comparable to those of the Wuhan-Hu-1-imprinted adult sera against the D614G variant (**Fig. 1C**).

### Neutralization specificities differ between sera from primary XBB* infections versus sera imprinted by early vaccination

To examine the specificity of the neutralizing antibody activity of the polyclonal sera in the different groups described above, we used pseudovirus-based deep mutational scanning (21,22) to measure how mutations in the XBB.1.5 spike affect neutralization by each serum.

We have previously described that for *Wuhan-Hu-1-imprinted and XBB* infected adult sera*, the major neutralization-escape mutations are in the RBD (21). To facilitate comparison to the new data generated in this study, we have replotted those prior measurements in **Fig. 2A**. New for this study, we also measured how mutations to the XBB.1.5 spike affected neutralization by the *Wuhan-Hu-1 imprinted and XBB* infected children sera*. Similarly to the previously published adult sera, neutralization by these imprinted children sera was most affected by RBD mutations (**Fig. 2A**, **Fig. S2**), with the key sites of escape similar between the imprinted adult and children sera. For both sets of sera, there are sites in the NTD where mutations reduce neutralization, but these mutations are at RBD-NTD interface sites (e.g., 132, 200, 234, 236; see structural visualizations at https://dms-vep.org/SARS-CoV-2_XBB.1.5_spike_DMS_infant_sera/structure.html) that have previously been shown to mediate indirect escape from RBD-directed antibodies by modulating the RBD’s up/down movement (21,27–29).

**Figure 2.**
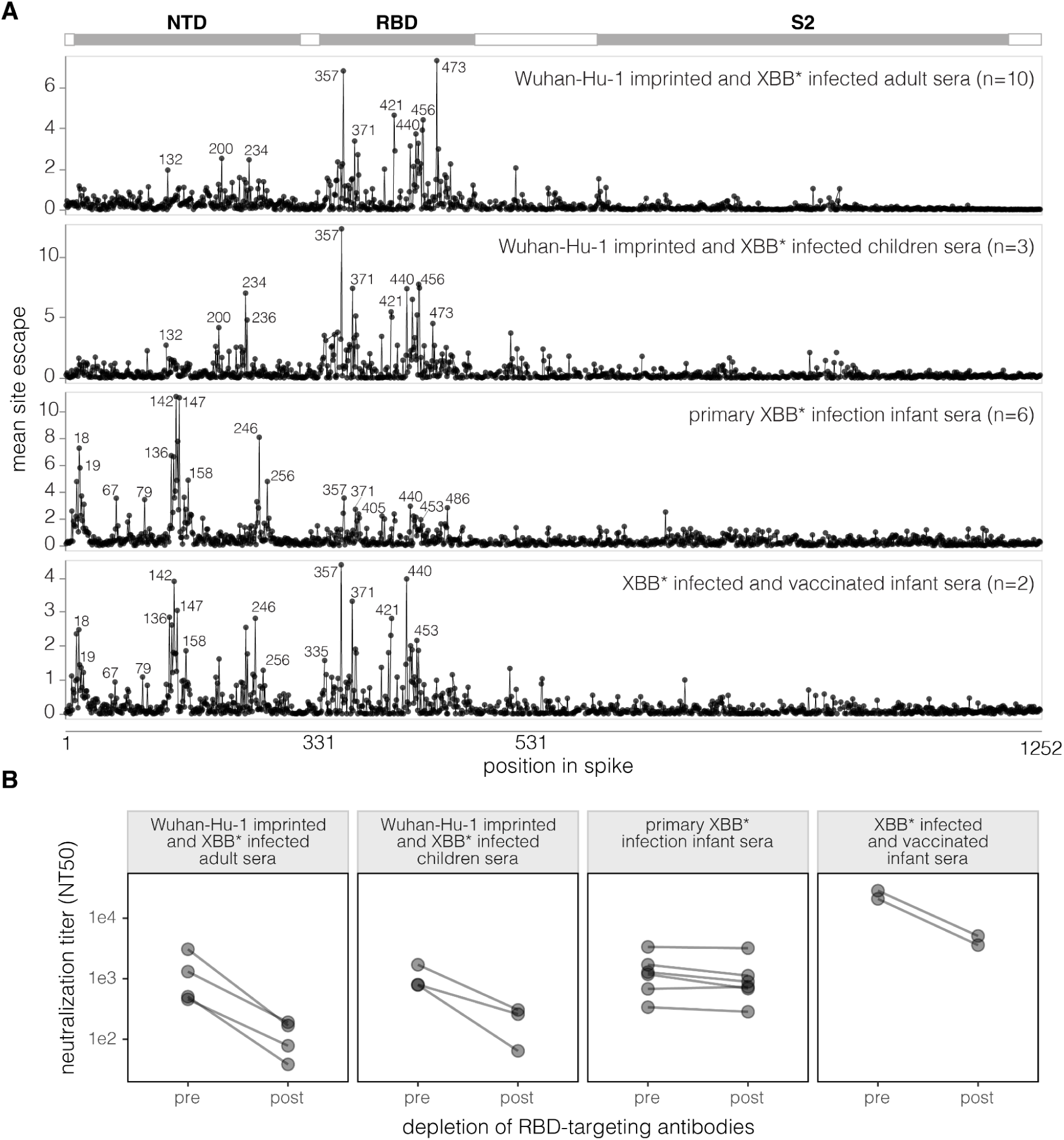
Neutralization specificities of sera from different exposure histories. **A.** Total neutralization escape caused by mutations at each site in the XBB.1.5 spike as measured by deep mutational scanning, averaged across the individual sera in each set. These plots show only mutations that have positive escape (reduced neutralization). See https://dms-vep.org/SARS-CoV-2_XBB.1.5_spike_DMS_infant_sera/average.html for an interactive version of this plot that makes it possible to zoom in on specific sites and examine the effects of individual mutations. See **Fig. S1** for escape plots for each individual serum (not averaged across the sera sets). **B.** Neutralization of pseudoviruses encoding the spike from XBB.1.5 by sera pre- and post-depletion of RBD-targeting antibodies depletion. For the *Wuhan-Hu-1 imprinted and XBB* infected adult sera*, these depletion assays were performed for four representative sera. See **Fig. S2** for details about the depletion assays.

In striking contrast to the imprinted adult and children sera, for the *primary XBB* infection infant sera*, the mutations that cause the greatest neutralization escape are in the NTD rather than the RBD (**Fig. 2A, Fig. S2**). Specifically, the top sites of escape are in the N1 (sites 14-28), N3 (sites 131-167), and N5 (sites 241-260) loops of the NTD, which together form the NTD antigenic supersite (18). These NTD sites are distal from the NTD-RBD interface and outside the region of the NTD where mutations affect the RBD’s up/down movement (21,27–29), indicating that the mutations in the NTD antigenic supersite are likely escaping NTD-targeting neutralizing antibodies directly rather than indirectly causing escape from RBD antibodies. The *primary XBB* infection infant sera* are also escaped a modest amount by RBD mutations, but for all but one of these sera the escape caused by NTD mutations substantially exceeds that caused by RBD mutations (**Fig. S2**).

However, secondary exposure of the infants to the XBB* spike via vaccination increases the focus of the neutralizing activity towards the RBD, as evidenced by the fact that the *XBB* infected and vaccinated infant sera* has major sites of escape in both the NTD and RBD (**Fig. 2A**, **Fig. S2**). While our study does not provide any direct insight into the mechanistic basis of this increased focus on the RBD after vaccination, we note that prior studies on adults have shown that vaccination after infection can lead to affinity maturation of potently neutralizing anti-RBD antibodies (30,31).

To validate the differences in the NTD-versus RBD-directed neutralizing activity measured in the deep mutational scanning, we depleted RBD-binding antibodies from individual sera (**Fig. S3**, **S4**) and measured the change in XBB.1.5 pseudovirus neutralization caused by the antibody depletion (**Fig. S3**, **S5**). Depletion of RBD-binding antibodies from *Wuhan-Hu-1-imprinted and XBB* infected adult sera* and *Wuhan-Hu-1-imprinted and XBB* infected children sera* greatly reduced neutralizing titers (**Fig. 2B**, **Fig. S5**), consistent with the deep mutational scanning showing that the major sites of escape from these sera are in the RBD. In contrast, depleting RBD-binding antibodies from *primary XBB* infection infant sera* had little effect on neutralizing titers (**Fig. 2B**, **Fig. S5**), again consistent with the deep mutational scanning showing that the major sites of escape from these sera are in the NTD rather than RBD. Depletion of RBD antibodies from the *Wuhan-Hu-1 imprinted and XBB* infected children sera* appreciably reduced neutralization titers (**Fig. 2B**, **Fig. S5**), consistent with the deep mutational scanning showing these sera are escaped by mutations in both the RBD and NTD.

### Spike mutations have different and even opposite effects on neutralization by sera imprinted with early vaccination versus XBB* infection

We examined how individual spike mutations affect neutralization at key sites for each of the different sera sets, considering both mutations that escape neutralization and those that increase neutralization (negative escape) (**Fig. 3A**, see also the interactive heatmaps at https://dms-vep.org/SARS-CoV-2_XBB.1.5_spike_DMS_infant_sera/average.html which have an option at the bottom to not floor escape at zero, and thereby see negative escape).

**Figure 3.**
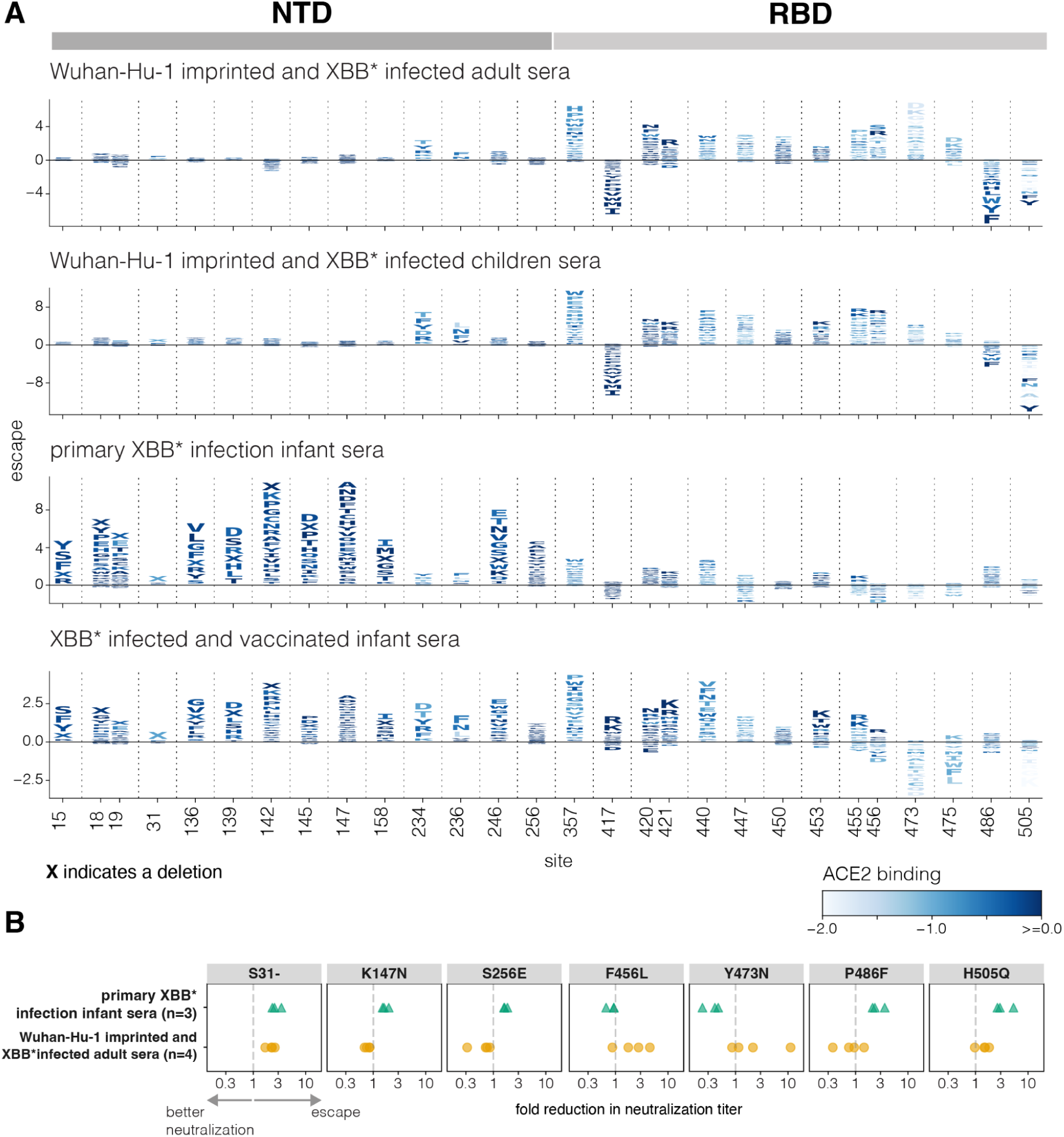
Effects of spike mutations on neutralization differ between sera imprinted by early vaccination versus infection with XBB* variants. **A.** Escape caused by individual mutations at key sites in spike for each sera set, with the escape values averaged across the individual sera in each set. The height of each letter is proportional to the escape caused by that mutation, with negative values (letters below the line at zero) indicating mutations that increase neutralization. Letters are coloured by their effects on full-spike ACE2 binding as previously measured in Dadonaite et. al. (2024) (21). The letter “X” indicates a single-residue deletion. See https://dms-vep.org/SARS-CoV-2_XBB.1.5_spike_DMS_infant_sera/average.html to interactively visualize all mutation-level escape values in heatmap form. **B.** Fold reduction in the neutralization titer against pseudoviruses expressing the indicated single mutant of the XBB.1.5 spike relative to the pseudovirus expressing the unmutated XBB.1.5 spike. Each point is the measurement for a different sera from that set (three sera were tested against each mutation for the *primary XBB* infection infant sera* and four sera were tested against each mutation for the *Wuhan-Hu-1 imprinted and XBB* infected adult sera*). The dashed line indicates no change in neutralization titer. Values greater than one indicate the mutation reduces neutralization and values below one indicate the mutation leads to better neutralization of the pseudovirus. The neutralization curves are in **Fig. S6**.

The *Wuhan-Hu-1 imprinted and XBB* infected adult* and *children* sera are similarly impacted by most mutations (top two panels of **Fig. 3A**), indicating that adults and children make neutralizing antibodies with similar specificities when they have similar exposure histories. The sites where mutations most strongly escape neutralization by these sera are nearly all in the RBD, and include sites 357, 420-421, 440, 447, 453, 455-456, 473 and 475. Mutations at some sites in the NTD (e.g., sites 234 and 236) also modestly escape these sera. These NTD escape sites are at the NTD-RBD structural interface and impact RBD up-down movement (21), and so mutations at these NTD sites probably indirectly escape RBD-targeting antibodies by putting the RBD in a more down conformation. There are also some sites in the RBD where mutations substantially increase neutralization (negative escape) by the imprinted adult and children sera, including sites 417, 486, and 505. Sites 417 and 486 of the Wuhan-Hu-1 spike were heavily targeted by neutralizing antibodies elicited by Wuhan-Hu-1 vaccination (16,32) but then acquired escape mutations during SARS-CoV-2 evolution, so mutating these sites in the context of the XBB.1.5 spike may restore neutralization by some Wuhan-Hu-1 elicited antibodies. Site 505 is targeted by neutralizing antibodies (15) but mutations at this site also put the RBD in a more up conformation (21) and enhance neutralization by antibodies targeting other RBD epitopes.

In contrast to the imprinted adult and children sera, the *primary XBB* infection infant sera* is most strongly escaped by mutations in the NTD (**Fig. 3A**), including at sites 15, 18-19, 136, 139, 142, 145, 147, 158, 246 and 256. These sites are all in the NTD antigenic supersite (18) which is on the exposed surface of the NTD in the full spike structure, and distal from the NTD-RBD interface where NTD mutations impact RBD up-down movement. Notably, NTD mutations that affect up-down movement of the RBD and escape the imprinted adult and children sera (e.g., at sites 234 and 236) have little impact on the *primary XBB* infection infant sera*, probably because the neutralizing activity of these sera is mainly due to antibodies targeted to the NTD rather than the RBD. Therefore, for the *primary XBB* infection infant sera*, NTD mutations are direct mediators of neutralization escape—whereas for the imprinted adult and children sera, NTD mutations mostly affect neutralization by indirectly escaping RBD antibodies via modulation of the RBD’s up/down conformation. Some RBD mutations at sites like 440 and 486 do modestly escape the *primary XBB* infection infant sera*, consistent with the fact these sera have some RBD-directed neutralization. However, mutations at many of the RBD sites that most strongly escape imprinted sera (e.g., 456, 473, 475) cause no escape from the *primary XBB* infection infant sera*.

The *XBB* infected and vaccinated infant sera* retains the NTD escape mutations of the *primary XBB* infection infant sera*, but also are more strongly affected by RBD mutations (**Fig. 3A**), as expected given the results in the prior section showing increased RBD-directed neutralizing activity for these sera (**Fig. 2**). However, there are differences in the sites in the RBD that escape these sera versus the imprinted adult and children sera. In particular, mutations at sites 456, 473, and 475 strongly escape the imprinted sera but do not escape the *XBB* infected and vaccinated infant sera*, consistent with recent work showing that these three sites are part of an epitope (the “A1 epitope”) that is primarily targeted by antibodies elicited by early vaccination then boosted by Omicron exposures (15). Interestingly, many mutations at sites 473 and 475 actually increase neutralization by the *XBB* infected and vaccinated infant sera* via a mechanism that remains unclear. Because the *XBB* infected and vaccinated infant sera* has more RBD-directed neutralizing activity than the sera from the same infants who have not been vaccinated, it is also escaped to a greater extent by mutations at NTD sites like 234 and 236 that put the RBD in a more down conformation.

To validate the highly divergent effects of mutations observed in the deep mutational scanning, we performed pseudovirus neutralization assays against the *Wuhan-Hu-1 imprinted and XBB* infected adult sera* and *primary XBB* infection infant sera* for seven different point mutants of the XBB.1.5 spike (**Fig. 3B**, **S6**). Consistent with the deep mutational scanning, the NTD mutations K147N and S256E escaped only the *primary XBB* infection infant sera*. Also consistent with the deep mutational scanning, RBD mutations F456L and Y473N escaped only the *Wuhan-Hu-1 imprinted and XBB* infected adult sera* whereas P486F reduced neutralization by the *primary XBB* infection infant sera* but enhanced neutralization by the *Wuhan-Hu-1 imprinted and XBB* infected adult sera*. H505Q appreciably reduced neutralization only by the *primary XBB* infection infant sera*. The only mutation we tested that clearly escaped neutralization by both sera sets was deletion of S31 in the NTD. This deletion likely directly escapes binding by some NTD antibodies, and also reduces serum neutralization by putting the RBD in a more down conformation (15,33,34). Note that deletion of S31 has occurred convergently in variants circulating in 2024 (35) and was responsible for the fitness advantage of the KP.3.1.1 variant relative to its parent KP.3.

### Primary infection by Wuhan-Hu-1 also elicits neutralizing activity with a substantial non-RBD-targeting component

We investigated if the high fraction of neutralizing activity directed towards the NTD for the *primary XBB* infection infant sera* is unique to infants infected by that variant, or is a more general feature of the response to primary SARS-CoV-2 infection. To do so, we examined primary *Wuhan-Hu-1 infection infant sera* and *primary Wuhan-Hu-1 infection adult sera*. These sera were collected from individuals who were infected by Wuhan-Hu-1 or a closely related variant early in the pandemic (**Table S1**).

To determine if neutralization by sera from primary Wuhan-Hu-1 infections is also dominated by non-RBD antibodies, we measured changes in D614G spike pseudovirus neutralization pre- and post-depletion of the RBD binding antibodies (**Fig. 4A, S7, S8**). Roughly half of the primary Wuhan-Hu-1 infection sera have a majority of their neutralizing activity from non-RBD targeting antibodies, although the percent of neutralization attributable to the RBD antibodies is generally greater for these primary Wuhan-Hu-1 infection sera than for primary XBB* variant infection sera (**Fig. 4**).

**Figure 4.**
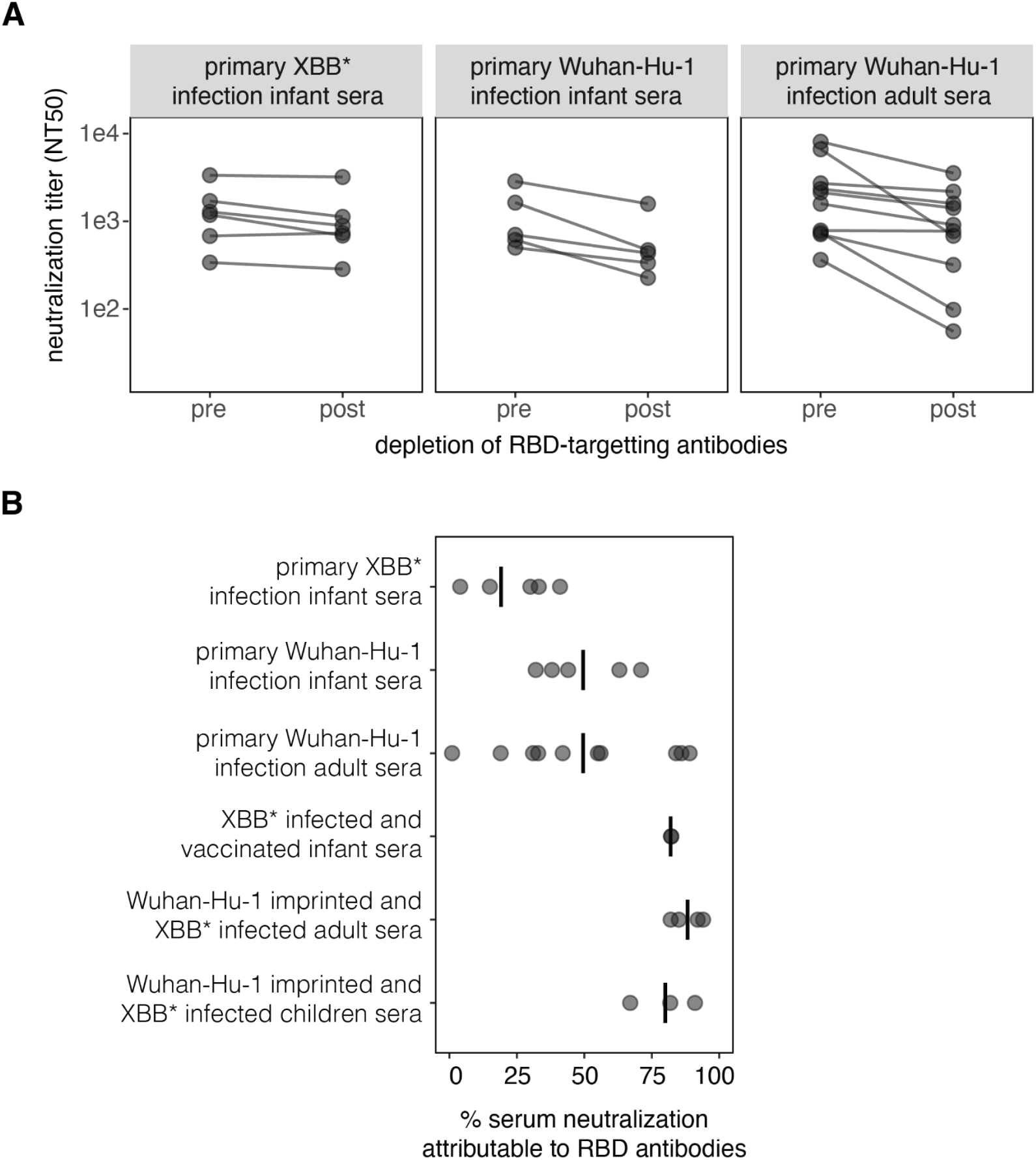
The neutralizing activity elicited by primary Wuhan-Hu-1 infection also has a substantial non-RBD targeting component. **A.** Neutralization of pseudoviruses expressing spike by sera pre- and post-depletion of RBD-targeting antibodies. The data for the *primary XBB* infection infant sera* are re-plotted from Fig. 2B and were generated using XBB.1.5 spike pseudoviruses and sera depleted with the XBB.1.5 RBD. The data for the *primary Wuhan-Hu-1 infection infant* and *adult sera* were generated using D614G pseudovirus and sera depleted with the Wuhan-Hu-1 RBD (note that D614G differs from Wuhan-Hu-1 only a by a single non-RBD mutation). **B.** Percent of serum neutralization attributable to RBD-binding antibodies for each serum set, calculated from the change in neutralization titers pre-versus post-depletion of RBD-binding antibodies. Each point is a different serum and the lines indicate the mean for each serum set.

The above results are in contrast to the past research by our group and others reporting that most sera from primary Wuhan-Hu-1 infections in adults have the majority of their neutralizing activity directed towards the RBD (12,13,36). We wondered if the differences could be due to technical aspects of how the neutralization assays were performed. Many SARS-CoV-2 studies perform neutralization assays using target cells that express high levels of ACE2, such as a 293T-ACE2 cell line generated early in the pandemic by our lab (37). However, we and others have since shown that target cells that express high levels of ACE2 tend to make neutralization assays less sensitive to antibodies that do not directly block RBD-ACE2 binding compared to target cells that express more modest levels of ACE2 (38,39). For the current study, all RBD-depletion neutralization assays were performed using a 293T cell line expressing medium levels of ACE2 (38). We compared neutralization titers of the *primary Wuhan-Hu-1 infection adult sera* pre- and post-depletion of RBD binding antibodies using medium-ACE2 cells versus the higher-ACE2 expressing 293T-ACE2 cells (38). Indeed, non-RBD antibodies were measured to contribute more to serum neutralization when the assays were performed using the cells expressing modest levels of ACE2 (**Fig. S9**). This finding underscores how the quantitative contribution of non-RBD antibodies to serum neutralization depends to some extent on the target cell line used for the measurements.

### Both NTD and RBD sites are highly variable during the evolution of SARS-CoV-2 in humans

We compared the natural diversity of the SARS-CoV-2 spike protein among human sequences to the sites of escape measured for both the primary infection infant sera and the imprinted adult sera (**Fig. 5A**). Natural diversity in spike is heavily concentrated in several regions of the NTD and throughout the RBD. The regions of greatest NTD diversity substantially overlap with the key sites of escape for the primary infection infant sera, whereas the regions of greatest RBD diversity substantially overlap with the key sites of escape for the imprinted adult sera (**Fig. 5A**).

**Figure 5.**
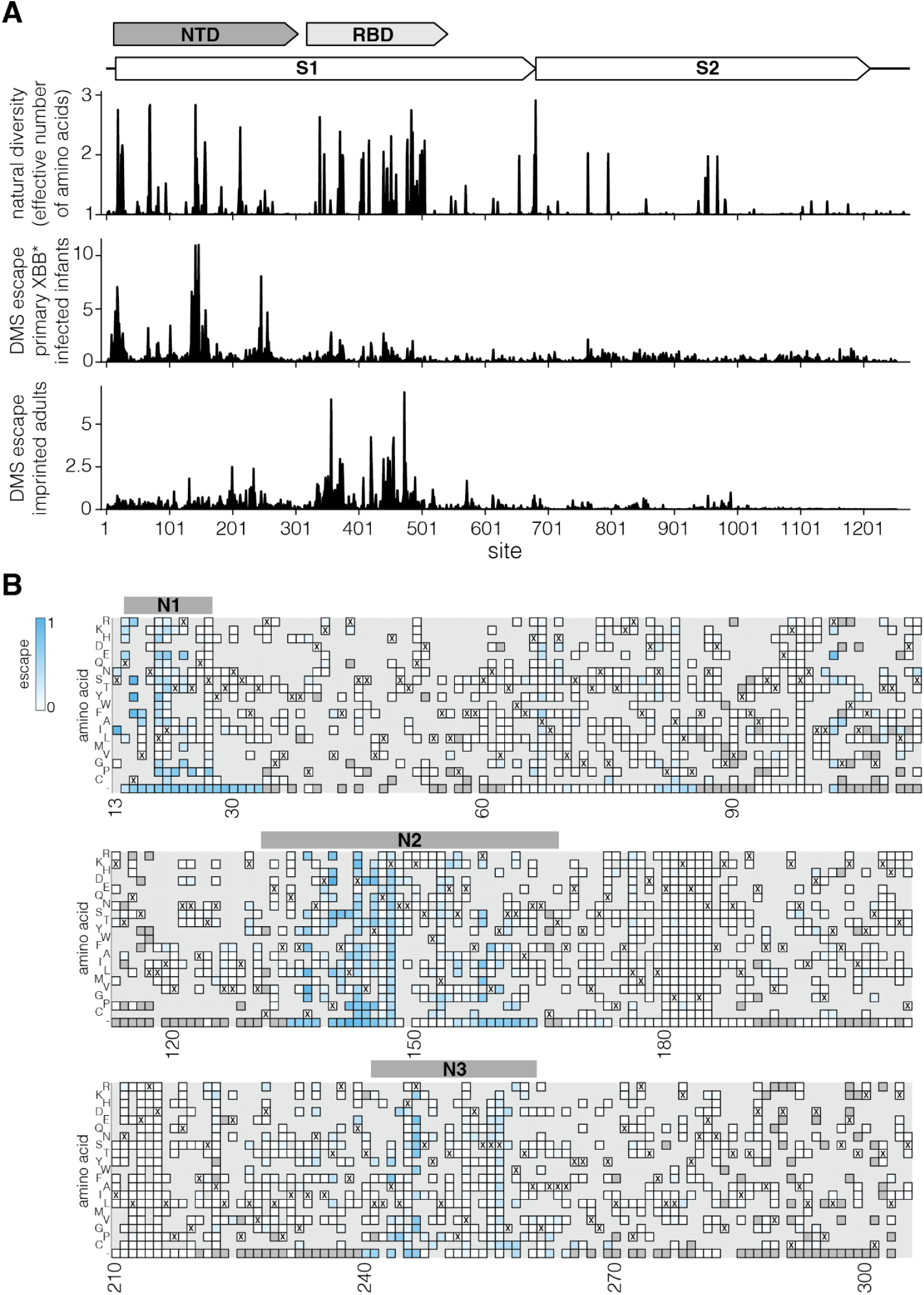
Sequence variability of SARS-CoV-2 spike compared to deep mutational scanning measured sera escape. **A.** A schematic of SARS-CoV-2 spike structure is at the top. The first plot below the schematic shows the amino-acid diversity in spike since 2020 calculated as the number of effective amino acids per site (exponential of the Shannon entropy), where one indicates a fully conserved site and values greater than one indicates more diversity. The second and third plots show the total escape at each site as measured in the deep mutational scanning for the *primary XBB* infant infection sera* and the *Wuhan-Hu-1 imprinted and XBB* infected adult sera* (these are the same data as in Fig. 2A). **B.** Effects of mutations in NTD on *primary XBB* infant infection sera* escape measured using deep mutational scanning. Mutations that lead to greater serum escape are colored blue. Mutations that are deleterious for cell entry are in dark grey. Mutations not in the library are in light grey. Bars above the heatmap show N1, N2 and N3 loops of the antigenic supersite. Only mutations that decrease serum neutralization are coloured. To also see mutations that increase serum neutralization, see the interactive heatmap at https://dms-vep.org/SARS-CoV-2_XBB.1.5_spike_DMS_infant_sera/average.html and toggle the floor escape at zero option at the bottom.

More specifically, the primary infection sera mostly targets the NTD antigenic supersite, composed of the N1 (sites 14-28), N3 (sites 131-167), and N5 (sites 241-260) loops in the NTD (18,20). The supersite has undergone extensive changes during the course of SARS-CoV-2 evolution in humans, including multiple amino acid mutations, deletions, and insertions (19,20). Many SARS-CoV-2 variants of concern had mutations in NTD regions where the deep mutational scanning shows strong escape for primary infection sera (40), including sites 19, 142-147, and 157-158 in the antigenic supersite. Deletions in the NTD loops that form the antigenic supersite were also common in major variants (19), and also lead to strong primary infection sera escape (**Fig. 5B**). Therefore, although the RBD remains the major target of neutralizing antibodies for imprinted adult sera (16,32,41), mutations in the NTD clearly also play an important role in the antigenic evolution of SARS-CoV-2 especially with respect to primary infection sera.

## Discussion

We have shown that there are dramatic differences in the specificity of polyclonal serum neutralizing activity between individuals who had a single primary SARS-CoV-2 infection versus those imprinted by vaccination with early variants. While prior work has shown that the epitope-specificity of neutralizing antibodies targeting the RBD differs depending on exposure history (6,15,42), our work shows that the differences extend to the domain level, with the neutralizing activity of primary XBB* infection infant sera mostly targeting the NTD rather than the RBD.

The reason that primary XBB* infection infant sera mostly targets the NTD whereas sera from imprinted individuals primarily targets the RBD remains unclear, but several lines of evidence suggest it relates to the exposure history rather than the age of the individuals. First, young children imprinted by early vaccination and then infected by XBB* have neutralizing specificities resembling those of similarly imprinted adults, suggesting young children develop a RBD-focused neutralizing response similar to adults when they have similar exposure histories. Second, both adults and children who have only been infected with an early Wuhan-Hu-1-like variant also tend to have serum neutralizing activities with substantial non-RBD (presumably NTD) targeting, albeit not to the same extent as the infants with primary XBB* infections. We also note that past research has suggested that mRNA vaccination with prefusion stabilized spike tends to elicit a more RBD-focused neutralizing response compared to infection (36).

Our results suggest that the neutralizing response to a single primary infection contains substantial NTD-targeting activity for multiple variants, but for infection with more recent XBB* variants, this NTD-targeting comprises the substantial majority of the neutralizing activity. Why XBB* primary infection elicits an even more NTD-targeted neutralizing response than primary infection by early variants remains unclear, but we note that recent variants like XBB* tend to have spikes with RBDs in a more down conformation, which hides some neutralizing RBD epitopes (43–45). In addition, more recent variants have mutated key neutralizing RBD epitopes present in early variants that are targeted by public near-germline antibody specificities (9,15,46), perhaps reducing the inherent immunogenicity of the RBD.

However, subsequent boosting by XBB.1.5 vaccination increases the RBD targeting of serum neutralization of infants initially infected by XBB* variants. We hypothesize that the increased RBD-directed neutralization after a secondary vaccination is due to affinity maturation of potently neutralizing anti-RBD antibodies (30,31). But whereas XBB* infected and then vaccinated infants have substantial neutralizing activity targeted to both the NTD and RBD, adults imprinted with early variants and then infected with XBB* have their neutralizing activity against XBB.1.5 directed overwhelmingly to the RBD. This may be due largely to evolution of the spike. While both the NTD and RBD have acquired numerous substitutions, the overall changes in the NTD have been more dramatic. The NTD has acquired multiple deletions, insertions, and substitutions that have largely obliterated cross-reactive epitopes in its antigenic supersite (19,20), whereas the RBD has evolved primarily by point mutations that have escaped many antibodies but led to little overall conformational change in the protein domain (6,13,47). For this reason, imprinted adults still produce cross-reactive antibodies targeting the RBDs of both early and recent variants (6,48,49), whereas NTD-specific antibodies provide limited cross-variant neutralization (41) and there is less boosting of cross-reactive neutralizing NTD antibodies upon exposure to new variants (50,51). The existence of more imprinted cross-reactive RBD than NTD antibodies likely leads to more RBD-targeted neutralization as individuals are repeatedly exposed to divergent variants.

Regardless of the mechanistic reason for more NTD-targeted neutralization in individuals with only primary infections, our results help explain the driving force behind the extensive antigenic changes in the NTD during the evolution of SARS-CoV-2 in humans (18–20). Notably, the recent evolution of SARS-CoV-2 has been dominated by mutations in the NTD, with an S31 deletion, T22N, F59S, K182R, Q183H, and R190S mutations convergently arising in multiple JN.1-descended variants (35). Some of these NTD mutations act in part by affecting RBD-directed neutralization by modulating the RBD’s up/down conformation (21,27–29), but many also directly affect neutralization by NTD-targeting antibodies (51). Because many studies of SARS-CoV-2 variant neutralization exclusively utilize sera from imprinted adults, these effects on NTD-directed neutralization may be largely overlooked. While the RBD remains the dominant target of neutralization by sera from most individuals (12–14), effects of both NTD and RBD mutations should be considered when it comes to their impact on evolution of new SARS-CoV-2. In particular, testing of sera from less imprinted individuals may be important to capture the effects of some NTD mutations.

Finally, our study demonstrates that mutations in the spike protein can have very different, and sometimes even opposing, effects on serum antibody neutralization depending on an individual’s exposure history. While the large majority of the global human population has been imprinted by early variants, a small but growing fraction of the population has been imprinted by more recent variants. Most of the individuals imprinted by more recent variants are young children, who may play an important role in shaping the evolution of respiratory viruses (52–54). As population heterogeneity in SARS-CoV-2 exposure histories increases, person-to-person variation in the antigenic effects of viral mutations is likely to influence the emergence of new SARS-CoV-2 variants and the susceptibility of different sub-populations, a phenomenon studied widely for influenza viruses (52,55–57).

## Supplementary Figures

**Supplementary Figure 1.**
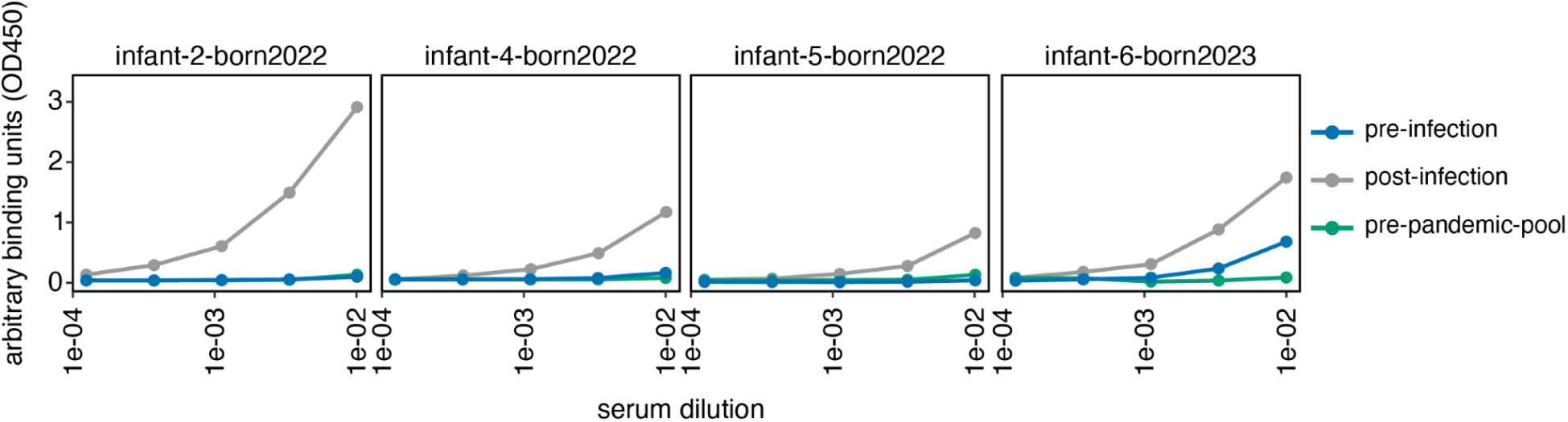
Binding of the infant sera to RBD-coated ELISA plates pre- and post-infection with a XBB* variant. Sera binding to XBB.1.5 RBD-coated plates measured using enzyme-linked immunosorbent assay (ELISA). The low RBD binding activity in the sera collected prior to infection indicates maternal SARS-CoV-2 antibodies had decayed to very low levels prior to infection. As a negative control, we binding is also shown for a pool of human sera collected prior to the beginning of the COVID-19 pandemic. Only four of the 6 infants in the *primary XBB* infection infant sera* group had pre-infection sera available. The pre-infection sera was collected 11-46 days before infection and therefore at the time of infection maternal antibodies may have decayed further. See **Table S1** for details on sera collection times.

**Supplementary Figure 2.**
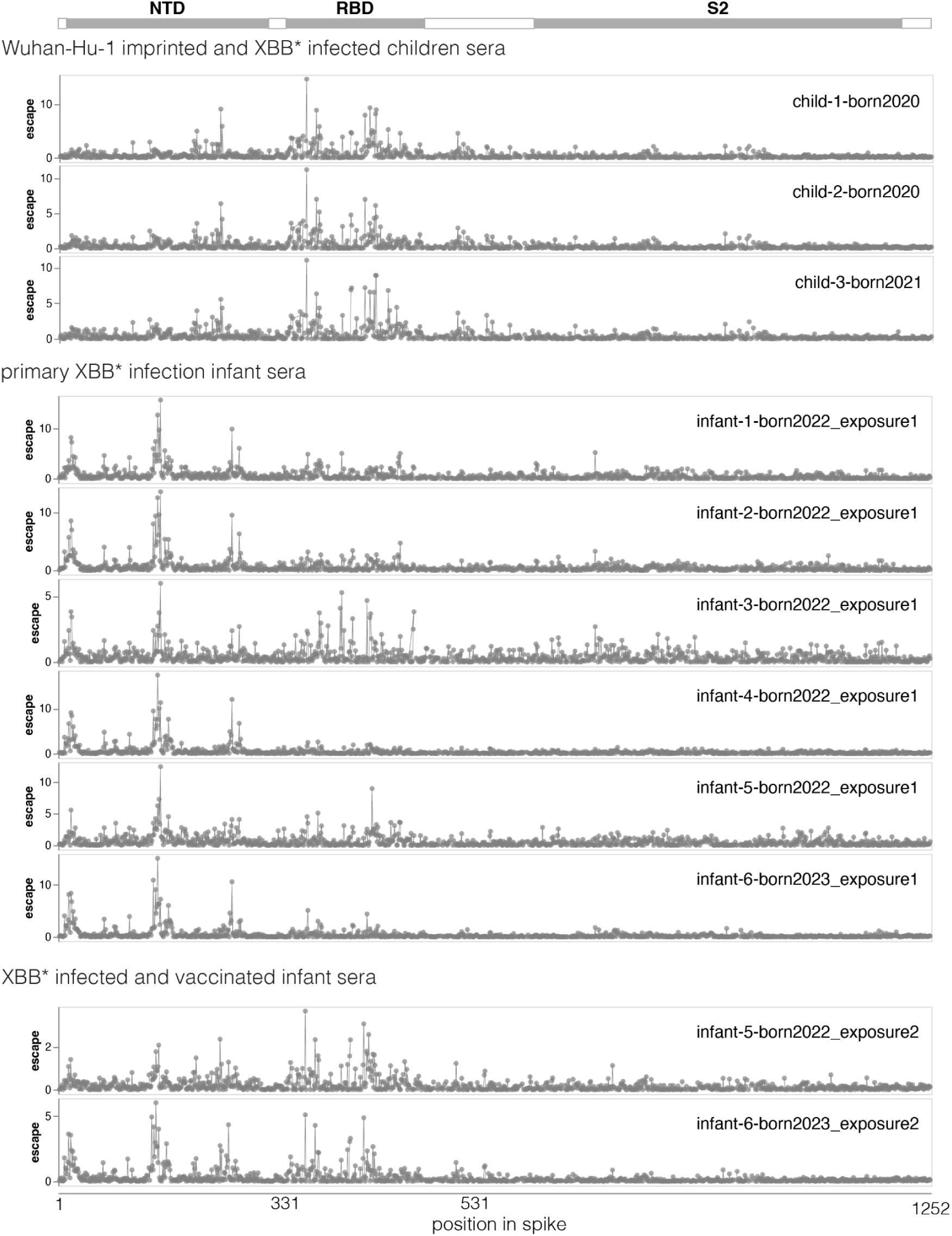
Deep mutational scanning escape maps for individual sera. These plots are similar to those in Fig. 2A except they show escape for the individual sera in each set rather than the average across all sera in each set. This figure only shows the plots for the three sera sets that were newly characterized in this study; the plots for the *Wuhan-Hu-1-imprinted and XBB* infected adult sera* have been previously published in Dadonaite et al. 2024 (21). See https://dms-vep.org/SARS-CoV-2_XBB.1.5_spike_DMS_infant_sera/individual.html for interactive versions of these plots for all four sera sets.

**Supplementary Figure 3.**
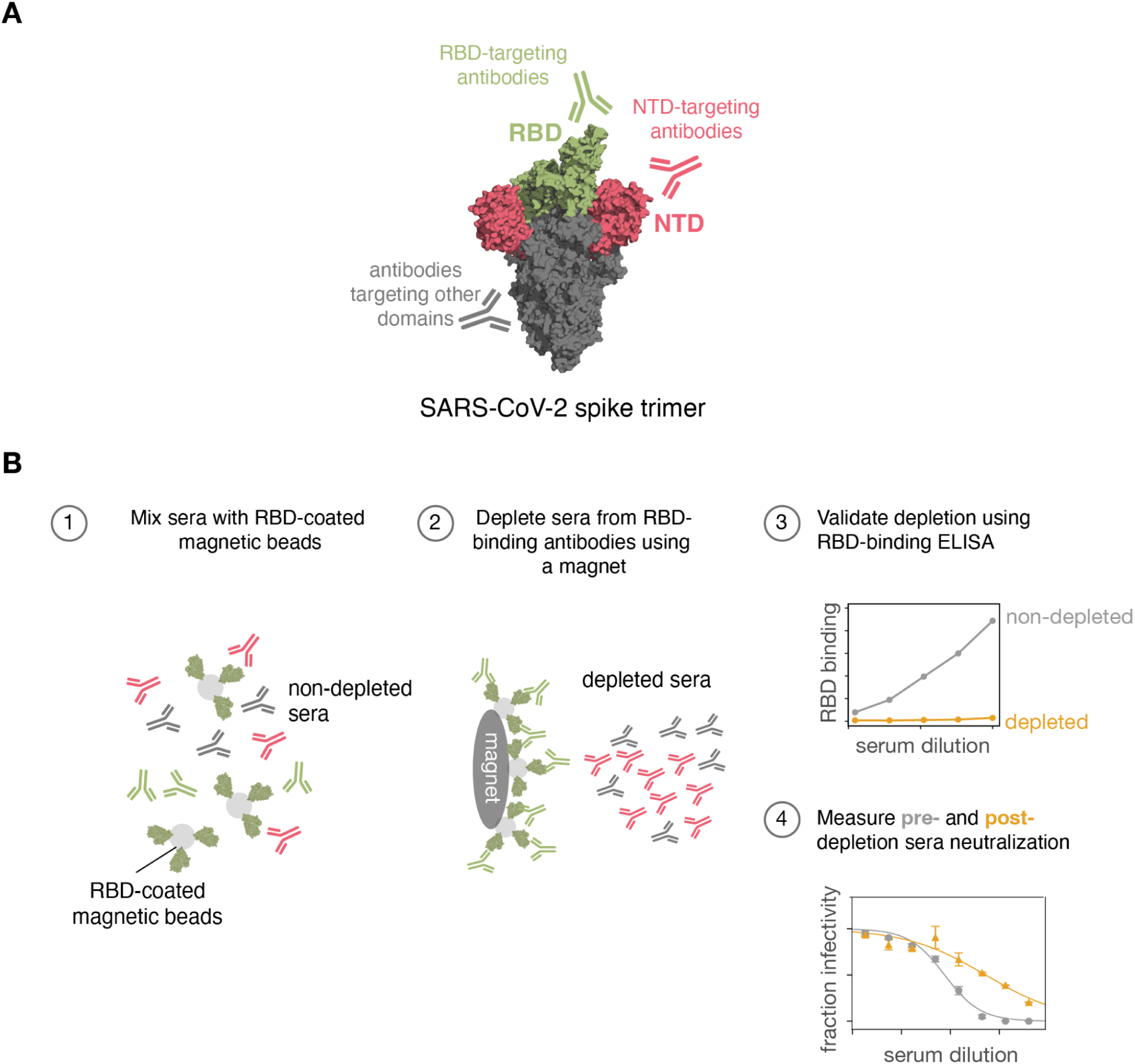
Depletion of RBD-binding antibodies from sera to measure their contribution to neutralization. **A.** Schematic of SARS-CoV-2 spike trimer labeling different domains (PDB: 6XM4). **B.** Each serum was mixed with RBD coated magnetic beads to allow RBD-targeting antibodies to bind the beads. After binding, a magnet was used to deplete sera of the beads. To validate successful RBD-binding antibody depletion enzyme-linked immunosorbent assay (ELISA) was used. Depleted and non-depleted sera was used to perform pseudovirus neutralization assays.

**Supplementary Figure 4.**
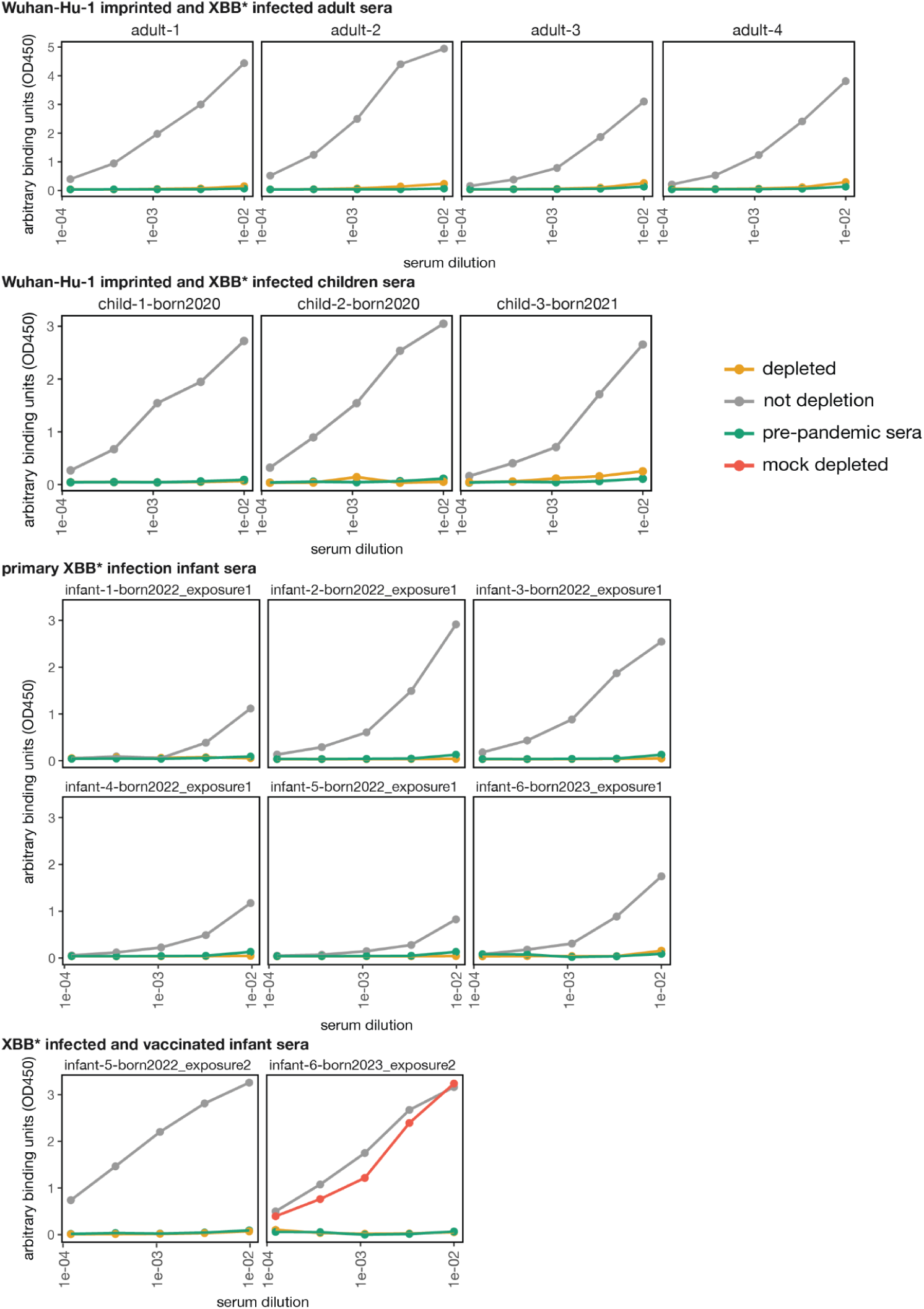
Binding of sera to RBD-coated ELISA plates pre- and post-depletion of RBD-binding antibodies. Depleted and non-depleted sera binding to XBB.1.5 RBD-coated plates measured using enzyme-linked immunosorbent assay (ELISA). Serum depletion was performed as described in **Fig. S2**. Sera was depleted by multiple rounds of XBB.1.5 RBD-coated bead incubation until its binding was similar to the pre-pandemic sera (sera collected prior to 2020 that should not contain any antibodies specific to SARS-CoV-2). A mock depleted sample, which was incubated with beads lacking XBB.1.5 RBD, is shown for *infant-6-born2020_exposure2* serum.

**Supplementary Figure 5.**
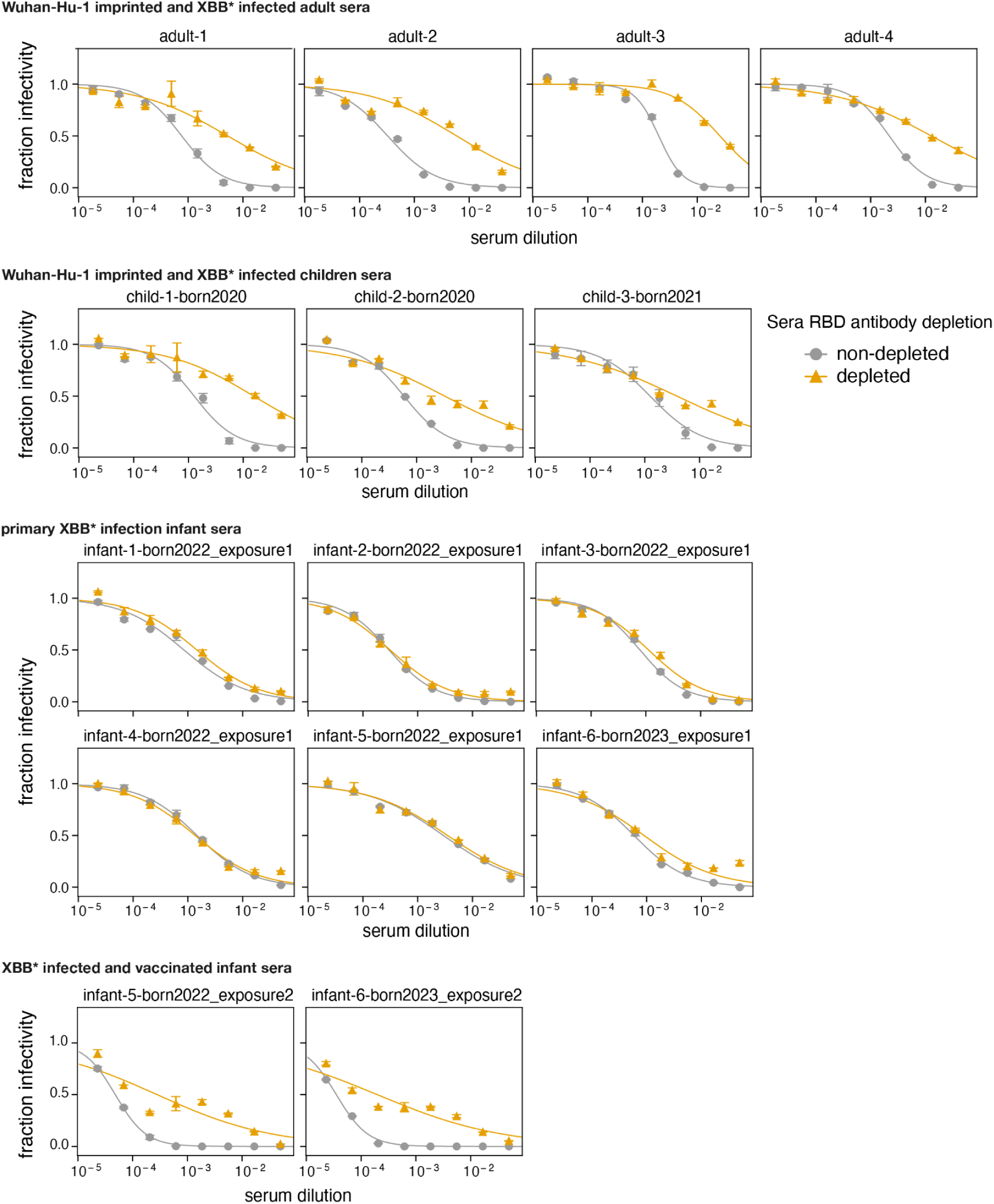
Neutralization of XBB.1.5 pseudovirus pre- and post-depletion of RBD-binding antibodies. Neutralization curves for XBB.1.5 pseudovirus against depleted and non-depleted sera from **Fig. S3**. Neutralization assays were performed on 293T cells expressing medium levels of ACE2 as described in Farrell et al. 2022 (20). These target cells enable better detection of the neutralizing activity NTD-binding antibodies than standard 293T-ACE2 cells that express higher levels of ACE2.

**Supplementary Figure 6.**
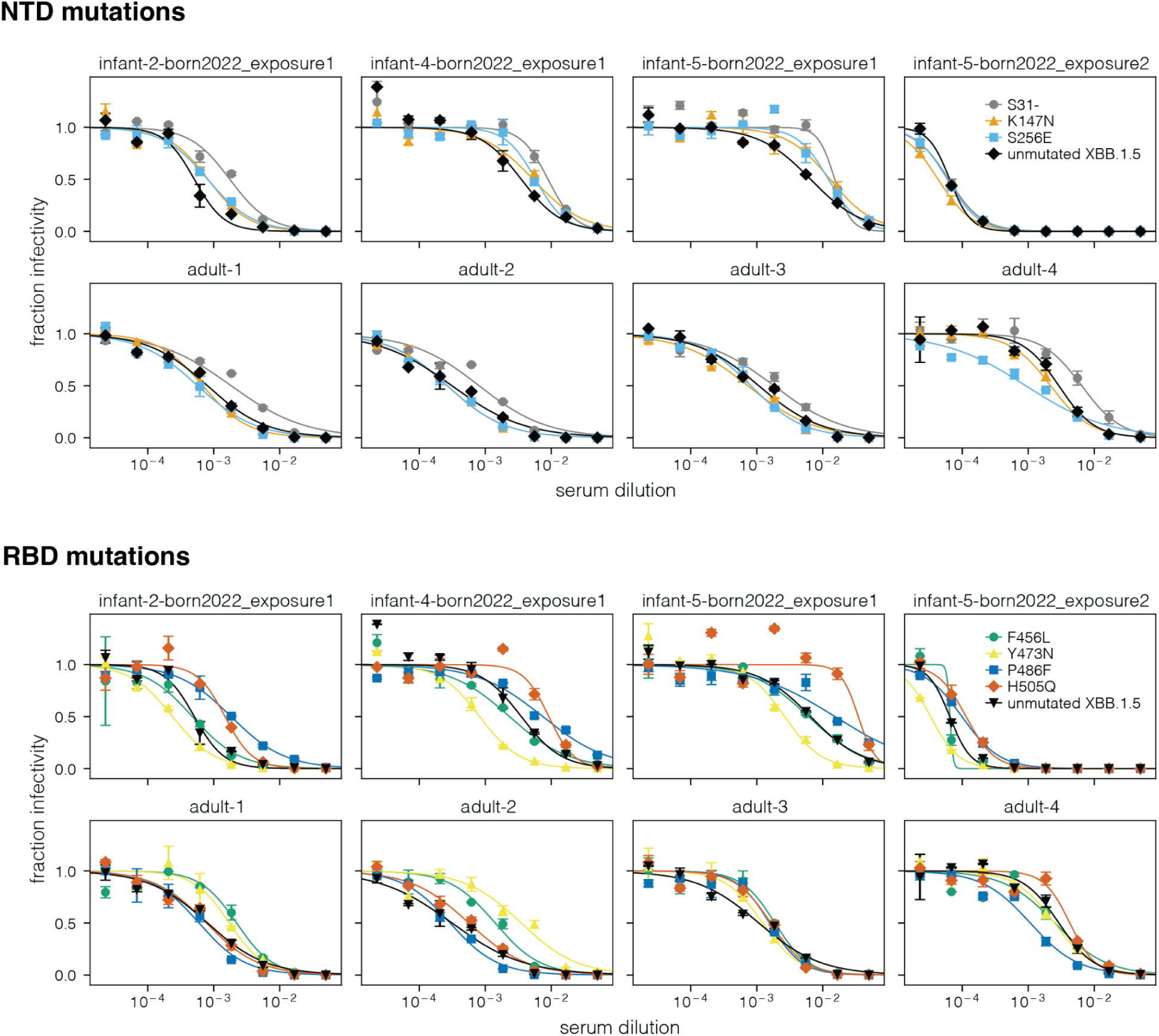
Pseudovirus neutralization of different XBB.1.5 spike mutants. Neutralization curves for unmutated XBB.1.5 and mutant XBB.1.5 spike pseudoviruses against different sera. All mutants contain a single amino-acid change on the background of the XBB.1.5 spike.

**Supplementary Figure 7.**
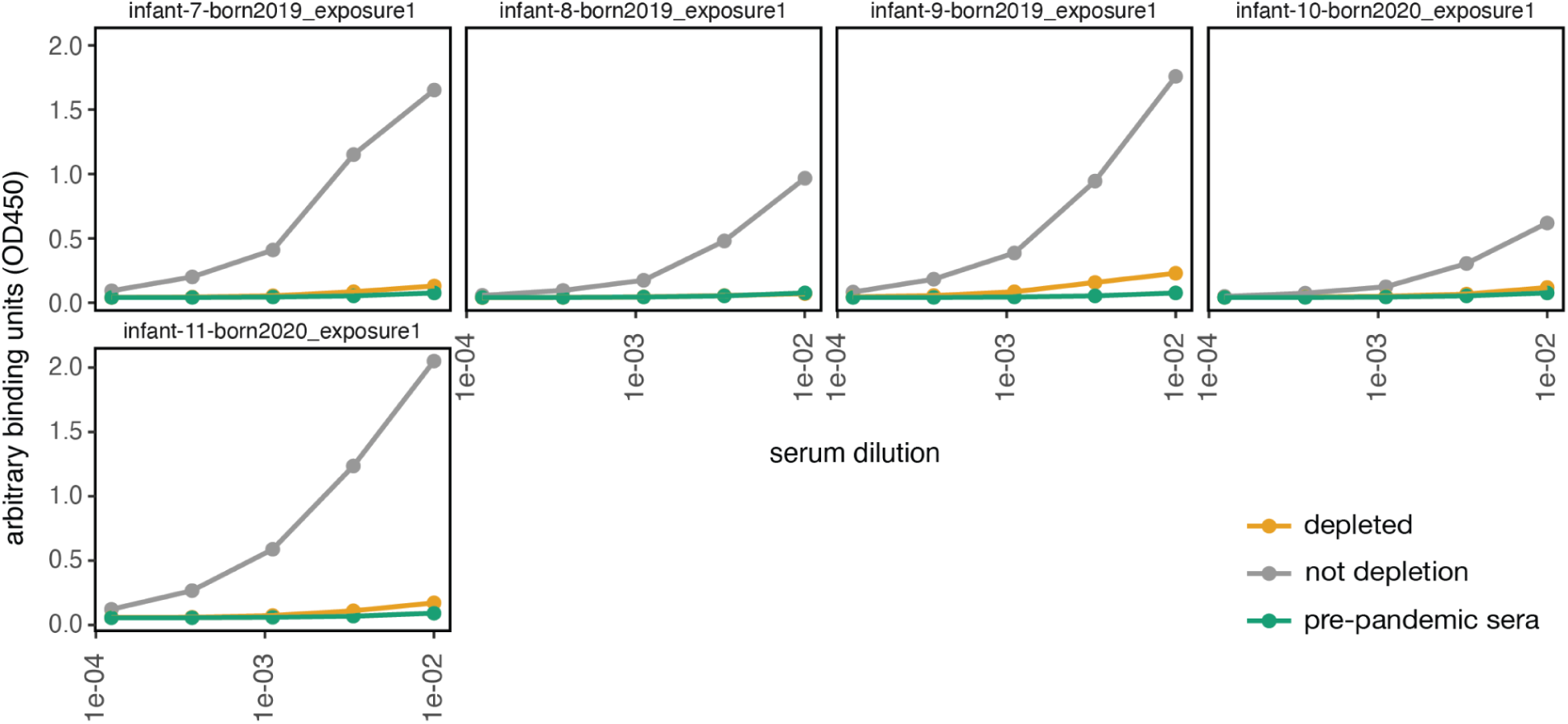
Binding of sera to RBD-coated ELISA plates pre- and post-depletion of RBD-binding antibodies for primary Wuhan-Hu-1 infant infection sera. Depleted and non-depleted sera binding to Wuhan-Hu-1 RBD-coated plates measured using enzyme-linked immunosorbent assay (ELISA). Serum depletion was performed as described in **Fig. S2** except using beads conjugated to the Wuhan-Hu-1 RBD rather than the XBB.1.5 RBD. Sera was depleted in multiple rounds of RBD-coated bead incubation until its binding was similar to the pre-pandemic sera (sera collected prior to 2020 that should not contain any antibodies specific to SARS-CoV-2). Note that depletions for *primary Wuhan-Hu-1 infection adult sera* were done for a previous study and that same pre- and post-depletion sera was reused for this study; pre- and post-depletion ELISA binding data for that sera comparable to that shown here can be found in the associated Greaney et al. (2021) paper (13).

**Supplementary Figure 8.**
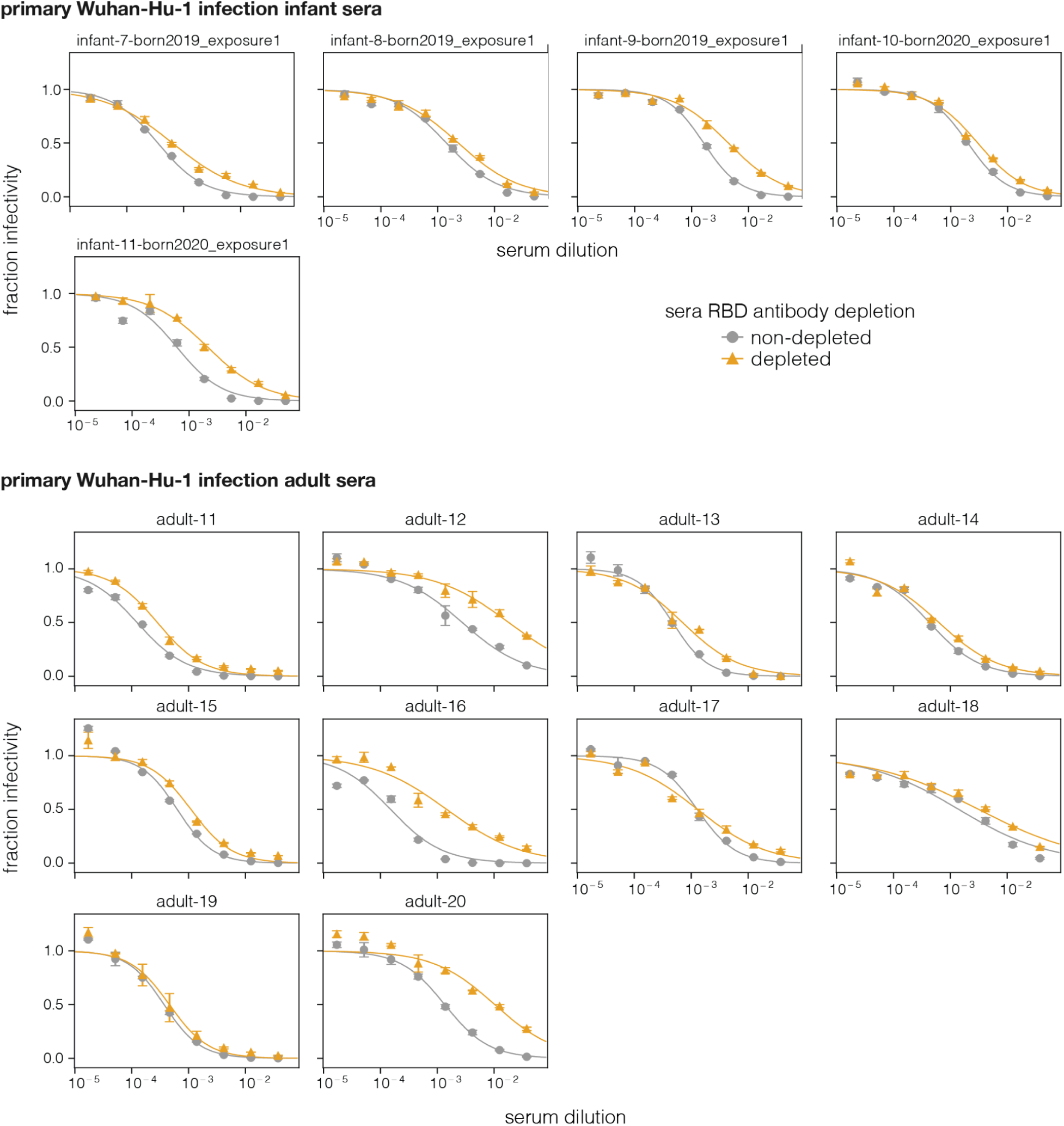
Neutralization of D614G pseudovirus pre- and post-depletion of RBD-binding antibodies for primary Wuhan-Hu-1 infections. Neutralization curves for D614G pseudovirus against depleted and non-depleted primary Wuhan-Hu-1 infection infant and adult sera. Neutralization assays were performed on 293T cells expressing medium levels of ACE2 cells as described in Farrell et al. 2022 (38).

**Supplementary Figure 9.**
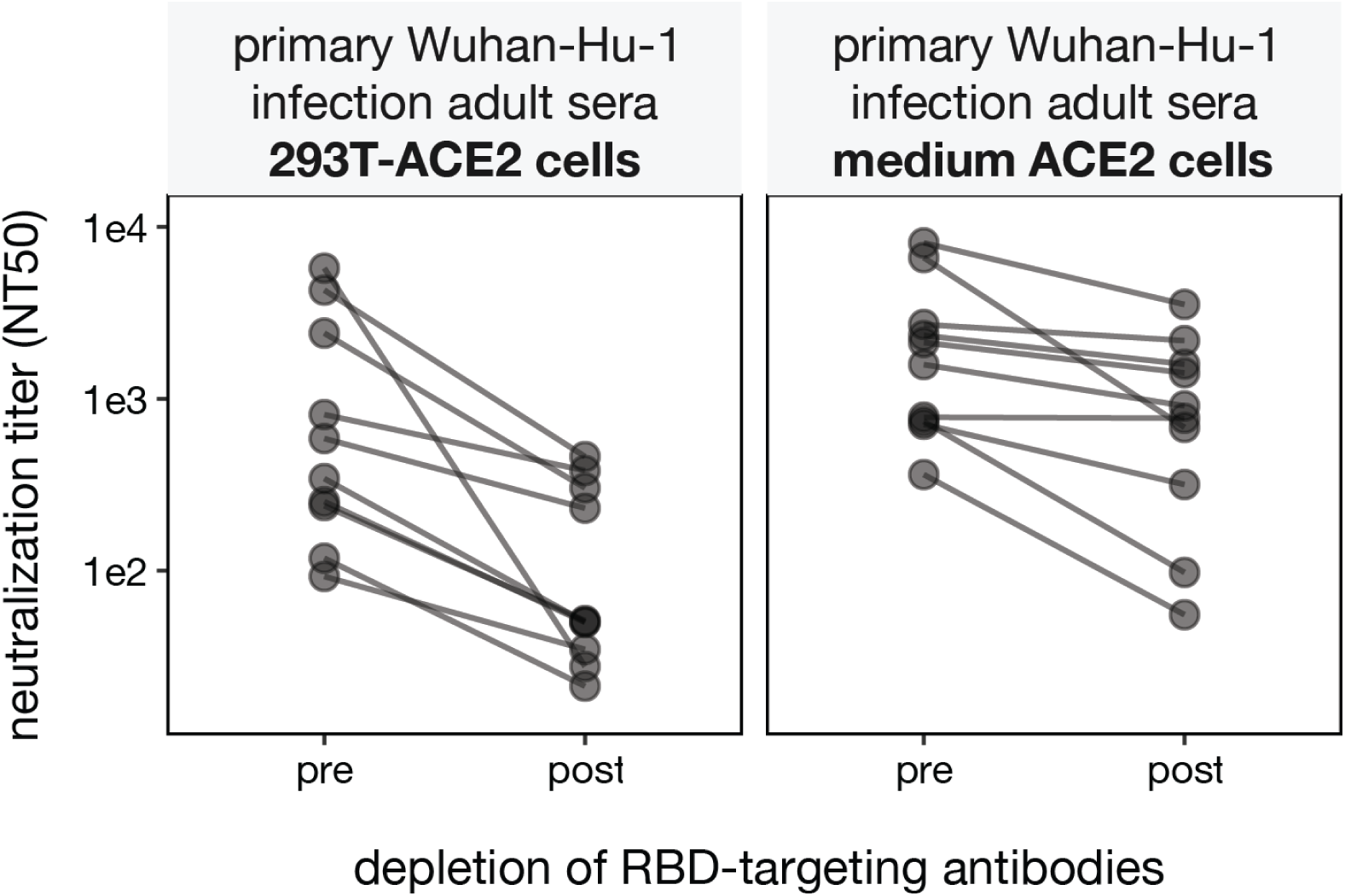
The extent that serum neutralization is due to RBD-binding antibodies depends on the ACE2 expression of the target cells. Neutralization of pseudovirus expressing the D614G spike by *primary Wuhan-Hu-1 infection adult sera* pre- and post-depletion of Wuhan-Hu-1 RBD targeting antibodies as measured on two different target cell lines. The left panel shows data replotted from Greaney et al. 2021 (13) for neutralization assays performed using the 293T-ACE2 cells (37), which express high levels of ACE2. The right panel shows new measurements made using the same pre- and post-depletion serum samples but with the 293T cells expressing medium levels of ACE2 (38). These medium ACE2 cells express less ACE2 than the 293T-ACE2 cells; see first supplementary figure of Farrell et al. 2022 (38).

## Supplementary Tables

https://docs.google.com/spreadsheets/d/1CfmnPS4G_eMFGGlSMzhalu7v0CDxQAgNAmP7YlPIRmk/edit?gid=0#gid=0

**Supplementary Table 1. Details of the individual sera used in this study**

Individual sera from each cohort used in this study. *Primary Wuhan-Hu-1 infected adult sera* and some of the *Wuhan-Hu-1 imprinted and XBB* infected adult sera* do not have sequencing data available for infections, however, based on infection dates the former cohort is most likely exposed to early strains closely related to Wuhan-Hu-1, and the latter to XBB*-related variants. The nasal swab from *infant-6-born2023_exposure1* did not yield sufficient coverage to confirm infecting variant but based on the date exposed this was most likely XBB* related variant. Negative values in the days post-exposure column indicate the number of days before exposure the sera were collected.

## Methods

### Data availability

We recommend readers visit https://dms-vep.org/SARS-CoV-2_XBB.1.5_spike_DMS_infant_sera/ for well-documented links to interactive visualizations and raw data. In addition:

● The full computational analysis pipeline is available on GitHub repository https://github.com/dms-vep/SARS-CoV-2_XBB.1.5_spike_DMS_infant_sera
●
● Interactive visualization comparing deep mutational scanning measured escape for the different sera sets used in this study is at https://dms-vep.org/SARS-CoV-2_XBB.1.5_spike_DMS_infant_sera/average.html
● Numerical data for the average serum escape for the different sera sets is at https://github.com/dms-vep/SARS-CoV-2_XBB.1.5_spike_DMS_infant_sera/blob/main/results/summaries/sera_group_avgs.csv
● Individual serum escape for *primary XBB* infected infant sera* is at https://github.com/dms-vep/SARS-CoV-2_XBB.1.5_spike_DMS_infant_sera/blob/main/results/summaries/primary_XBB_infection_infant_sera_per_antibody_escape.csv
● Individual serum escape for *XBB* infected and vaccinated infant sera* is at https://github.com/dms-vep/SARS-CoV-2_XBB.1.5_spike_DMS_infant_sera/blob/main/results/summaries/XBB_infected_and_vaccinated_infant_sera_per_antibody_escape.csv
● Individual serum escape for *Wuhan-Hu-1 imprinted and XBB* infected children sera* is at https://github.com/dms-vep/SARS-CoV-2_XBB.1.5_spike_DMS_infant_sera/blob/main/results/summaries/WuhanHu1_imprinted_and_XBB_infected_children_sera_per_antibody_escape.csv
● Individual serum escape for *Wuhan-Hu-1 imprinted and XBB* infected adult sera* is at https://raw.githubusercontent.com/dms-vep/SARS-CoV-2_XBB.1.5_spike_DMS_infant_sera/refs/heads/main/results/summaries/WuhanHu1_imprinted_and_XBB_infected_adult_sera_per_antibody_escape.csv. Note this dataset has been published before in Dadonaite et al. (2024).
● Raw sequencing data is available a BioProject PRJNA1210606

### Ethics statement

Wuhan-Hu-1 vaccinated and XBB* infected adult sera was collected with informed consent from participants in Hospitalized or Ambulatory Adults with Respiratory Viral Infections (HAARVI) study run by Washington University and approved by University of Washington Institutional Review Board (protocol number STUDY00000959). Primary Wuhan-Hu-1 infection sera was collected as part of longitudinal cohort study in Seattle February and July 2020 and has been described previously in Crawford et al. (2020) (58). Sera from infants and children was collected from participants of the IMPRINT Influenza Cohort, a longitudinal birth cohort being conducted in Cincinnati, Ohio. Mothers provide written informed consent for themselves and their child at enrollment in the third trimester of pregnancy. The study received approval from the Institutional Review Boards at Cincinnati Children’s Hospital Medical Center (2019-0629) and the University of Cincinnati Medical Center (SITE00000489).

### Sera samples

See **Supplementary Table 1** for all available metadata for the sera samples used in this study. Pre-pandemic sera used as a negative control for the RBD ELISAs was collected between 2017-2018 and came from Gemini Bio (100-110-100). All sera was inactivated at 56°C for 1 h and stored at −80°C long term.

### SARS-CoV-2 variant sequencing for infected infants

Nasal swab samples underwent nucleic acid extraction using the QIAGEN Viral RNA Mini Kit (Qiagen, Inc) on the QIAGEN QIAcube Connect instrument following the manufacturer’s recommendations. RNA samples were subjected to overlapping amplicon sequencing using the modified ARTIC3 protocol ((https://artic.network/ncov-2019) with the addition of primer booster sets as implemented in the Qiagen QiaseqDirect protocol. In brief, RNA was subjected to random primed cDNA synthesis followed by amplification in two pools of multiplexed primer sets resulting in overlapping amplicons spanning the entire genome. Subsequently, 24 cycles of polymerase chain reaction were utilized to add dual index primers and amplify SARS-Cov-2 amplicons. DNA concentrations were normalized, samples were pooled and then subjected to sequencing to a depth of at least 500,000 reads per sample utilizing paired 150 nucleotide reads on an Illumina NextSeq 500 sequencing machine (Illumina, Inc).

Raw sequence data were demultiplexed and then aligned against the ancestral Wuhan-1 genome (Accession MN908947) (59)using bwa-mem (60). Samtools commands “sort”, “index”, “view”, and “mpileup” (61) were applied sequentially, and the ivar “consensus” command (62) was used to output the consensus genome sequence. Variant identification and lineage calling were performed with the software Phylogenetic Assignment of Named Global Outbreak Lineages (Pangolin) version 4.3.1 (https://cov-lineages.org/resources/pangolin.html) (63).

### Deep mutational scanning library production

The XBB.1.5 full spike pseudovirus deep mutational scanning libraries used in this study were described previously (21). These libraries consist of spike pseudotyped in lentiviral particles. The libraries were designed to include all accessible and tolerated amino-acid mutations in spike; see (21) for more details. The pseudoviruses used for the deep mutational scanning were generated via the process described in (21). The only change in library virus generation was an added ultracentrifugation step to further increase library virus titers. To do so, virus-containing supernatants were concentrated by ultracentrifugation over 20% sucrose cushion (made in buffer containing 100 mM NaCl, 20 mM Tris-HCl (pH 7.4), 5 mM CaCl_2_) by spinning at 100,000g for 1 hour using Beckman Coulter SW 32 Ti rotor. Virus pellet was resuspended in a resuspension buffer (10 mM Tris-HCl (pH 7.4), 100 mM NaCl, 0.1 mM EDTA).

### Serum escape selections using deep mutational scanning libraries

Serum selections were performed as described previously (21). In brief, serum neutralization was first determined using a standard pseudovirus neutralization assay (described below). Concentrated deep mutational scanning library viruses were mixed with RDPro pseudovirus at 1-2% of total library transcription units used in an experiment. RDPro is used as a non-neutralizable standard in these experiments, which allows us to get quantitative escape values for each mutation in the library (21,22). Production of the non-neutralizable RDPro standard has been described previously (21). One-million transcription units of library virus (which covered the number of barcoded variants in the library variant at least 10-fold) were incubated with increasing serum concentrations for 45 min at 37°C. Starting serum concentrations used in these experiments were usually at IC98 as determined by a standard pseudovirus neutralization assay, and 3-fold and 6-fold greater (note that the actual neutralization of the libraries did not always precisely match the standard pseudovirus neutralization assay). Depending on serum availability and experimental results these concentrations needed to be adjusted for some sera. Generally, for good data fitting we aimed to use such serum concentrations where, when possible, at least some of the samples in the final analysis would have more than 90% of the variants in the library neutralized. 293T-ACE2 cells (37) (which express high amounts of ACE2) were then infected with these libraries. 12 hours after infection non-integrated virus genomes were recovered and amplicon libraries were prepared for multiplexed Illumina sequencing as described before (22). Illumina sequencing was performed on the NextSeq 2000 machine using P2 kit. For all sera experiments with two biological library replicates were performed.

### Analysis of serum escape mutations for deep mutational scanning experiments

To identify mutations in the deep mutational scanning library that affect serum neutralization *Polyclonal* software (v.6.12) (64), which is implemented in *dms-vep-pipeline-3* (https://github.com/dms-vep/dms-vep-pipeline-3), was used. *Polyclonal* uses a biophysical model to fit neutralization curves to varian-level infectivities and get mutation-level effects on serum neutralization. An example of *polyclonal* data fitting can be found at https://dms-vep.org/SARS-CoV-2_XBB.1.5_spike_DMS_infant_sera/notebooks/fit_escape_antibody_escape_Lib1-240903-infant-6-born2023_exposure2.html and the appendix to the interactive data visualization at https://dms-vep.org/SARS-CoV-2_XBB.1.5_spike_DMS_infant_sera/ provides links to similar notebooks for all experiments

### RBD-binding antibody depletion from sera

Streptavidin coated magnetic beads (Acrobio, SMB-B01) were washed 3 times in PBS using a magnetic stand. 100 µg of biotinylated XBB.1.5 RBD (Acrobio, SPD-C82Q3) was mixed with 1 mg of washed beads and incubated at room temperature for 1 hour. After incubation beads were spun dawn, placed on a magnet and washed 5 times with PBS+0.1% bovine serum albumin (PBS-B). After washing beads were resuspended at 1 mg/ml in PBS-B and stored at −80°C until further use. Wuhan-Hu-1 RBD was ordered already conjugated to magnetic beads (Acrobio, SPD-C52H3).

To deplete sera from RBD-binding antibodies, 70 µl of RBD-coated beads were mixed with 35 µl of sera and incubated on a rotating platform for 1 hour. Samples were spun down and placed on a magnet, supernatant was transferred to a new tube with a fresh aliquot of RBD-conjugated beads. This process was repeated multiple times until serum binding to RBD-coated plates by ELISA (see below) was no greater than the binding of sera collected pre-pandemic (in 2017-2018). For primary Wuhan-Hu-1 adult and infant sera Wuhan-Hu-1 conjugated beads were used to perform depletions, for all other sera XBB.1.5 conjugated beads were used.

### Enzyme-linked immunosorbent assay

To validate the depletion of RBD-binding antibodies from sera, Immunlon 2HB plates 96-well plates (Thermo Scientific 3455) were coated with 50 µl of the RBD protein at 0.5 µg/mL in PBS at 4°C overnight. For primary Wuhan-Hu-1 adult and infant sera plates were coated with Wuhan-Hu-1 RBD, for all other sera XBB.1.5 RBD was used. Next day the unbound RBD was removed by washing the plates 3 times with PBS + 0.1% Tween-20 (PBS-T). Plates were blocked by incubation with 150 uL of 3% (w/v) dried milk in PBS-T for 1 hour at room temperature. In a separate plate serial dilutions of depleted, non depleted sera and pre-pandemic sera were prepared in 1% (w/v) dried milk in PBS-T. Serial sera dilutions were transferred on blocked ELISA plates and incubated for 1 hour at room temperature. Afterwards, plates were washed 3 times with PBS-T and 50 µl of 1:3000 diluted horseradish peroxidase conjugated goat anti-human IgG-Fc antibody (Bethyl Labs, A80-104P) was added and incubated at room temperature for 1 hour. Subsequently, plates were washed 3 times with PBS-T and 100 µl of horseradish peroxidase substrate (Millipore, ES001-500ML) was added. Reaction was allowed to proceed for 5 minutes before 100 µl of 1N HCl was added. Light absorption was detected by reading plates at 450 nm within 5 min of adding HCl.

### Pseudovirus neutralization assays

Pseudovirus neutralization assays were performed as described previously (37) with the following modifications. To produce pseudoviruses 293T cells were transfected with Gag/pol expression plasmid (BEI: NR-52517), pHAGE6_Luciferase_IRES_ZsGreen luciferase-containing lentivirus backbone and spike expression plasmid. All wild-type and mutated spike expression plasmids used in this study can be found at https://github.com/dms-vep/SARS-CoV-2_XBB.1.5_spike_DMS_infant_sera/tree/main/validation_notebooks/plasmid_maps. 48 hours after transfection pseudovirus-containing cell supernatants were collected, filtered to remove cell debris, and titrated as described previously (37).

To perform neutralization assays 50,000 293T-ACE2 (37) or medium-ACE2 (38) cells were plated in the presence of 2.5 µg/ml of amphotericin B in black-walled poly-L lysine coated 96-well plates. For neutralization assays with RBD-binding antibody depleted or pre-depleted sera medium-ACE2 cells were used, for all other neutralization assays 293T-ACE2 cells were used. Amphotericin B has been shown previously to increase SARS-CoV-2 pseudovirus titers (22). For neutralization assay starting serum dilution was 1:20 and it was serially diluted 1:3 in a 96 well plate. Note that in the process of depleting RBD-binding antibodies sera was diluted 1:3, we accounted for this dilution during neutralization assays by using 3 times more of the depleted sera. Serum dilutions were mixed with pseudoviruses, incubated at 37°C for 45 min and then transferred to pre-plated cells. Relative light units in each well were detected 48 hours later using the Bright-Glo Luciferase Assay System (Promega, E2610).

### Cell lines

All cell lines were grown in D10 media (Dulbecco’s Modified Eagle Medium with 10% heat-inactivated fetal bovine serum, 2 mM l-glutamine, 100 U/mL penicillin, and 100 μg/mL streptomycin). Medium-ACE2 cells were additionally supplemented with 2 µg/ml of doxycycline. Cell-stored deep mutational scanning libraries were grown in D10 media using doxycycline-free FBS.

## Acknowledgments

This research was funded by the following grants from the NIAID/NIH to JDB: P01AI167966, the SAVES program (contract 75N93021C00015, option 18.C), and R01AI141707. JDB is an investigator at the Howard Hughes Medical Institute. This research was also supported by the Genomics & Bioinformatics Shared Resource, RRID:SCR_022606, awarded to the Fred Hutch/University of Washington/Seattle Children’s Cancer Consortium (P30 CA015704). The IMPRINT cohort is funded by NIH grant U01 AI144673-01 (Principal Investigator M.A.S.) and Open Philanthropy/Good Ventures. The Center for Clinical & Translational Science & Training at the University of Cincinnati provides support for IMPRINT database management and study conduct and is funded by the NIH Clinical and Translational Science Award program, grant UL1TR001425.

## Competing interests

J.D.B. and B.D. are inventors on Fred Hutch licensed patents related to the pseudovirus deep mutational scanning technique used in this paper. J.D.B. consults for Apriori Bio, Invivyd, Pfizer, the Vaccine Company, Moderna, and GSK. B.D. consults for Moderna. HYC has consulted for Bill and Melinda Gates Foundation and Ellume, and has served on advisory boards for Vir, Merck, Roche and Abbvie. She has received research funding from Gates Ventures. M.A.S. has received research funding from Cepheid and Pfizer and has consulted for Merck.

## Notes

https://dms-vep.org/SARS-CoV-2_XBB.1.5_spike_DMS_infant_sera/

https://github.com/dms-vep/SARS-CoV-2_XBB.1.5_spike_DMS_infant_sera

